# Tracking SARS-CoV-2 Omicron diverse spike gene mutations identifies multiple inter-variant recombination events

**DOI:** 10.1101/2022.03.13.484129

**Authors:** Junxian Ou, Wendong Lan, Xiaowei Wu, Tie Zhao, Biyan Duan, Peipei Yang, Yi Ren, Lulu Quan, Wei Zhao, Donald Seto, James Chodosh, Jianguo Wu, Qiwei Zhang

**Author notes:** Correspondence: Q.Z., J.W.,.

## Abstract

The current pandemic of COVID-19 is fueled by more infectious emergent Omicron variants. Ongoing concerns of emergent variants include possible recombinants, as genome recombination is an important evolutionary mechanism for the emergence and re-emergence of human viral pathogens. Although recombination events among SARS-CoV-1 and MERS-CoV were well-documented, it has been difficult to detect the recombination signatures in SARS-CoV-2 variants due to their high degree of sequence similarity. In this study, we identified diverse recombination events between two Omicron major subvariants (BA.1 and BA.2) and other variants of concern (VOCs) and variants of interest (VOIs), suggesting that co-infection and subsequent genome recombination play important roles in the ongoing evolution of SARS-CoV-2. Through scanning high-quality completed Omicron spike gene sequences, eighteen core mutations of BA.1 variants (frequency >99%) were identified (eight in NTD, five near the S1/S2 cleavage site, and five in S2). BA.2 variants share three additional amino acid deletions with the Alpha variants. BA.1 subvariants share nine common amino acid mutations (three more than BA.2) in the spike protein with most VOCs, suggesting a possible recombination origin of Omicron from these VOCs. There are three more Alpha-related mutations (del69-70, del144) in BA.1 than BA.2, and therefore BA.1 may be phylogenetically closer to the Alpha variant. Revertant mutations are found in some dominant mutations (frequency >95%) in the BA.1 subvariant. Most notably, multiple additional amino acid mutations in the Delta spike protein were also identified in the recently emerged Omicron isolates, which implied possible recombination events occurred between the Omicron and Delta variants during the on-going pandemic. Monitoring the evolving SARS-CoV-2 genomes especially for recombination is critically important for recognition of abrupt changes to viral attributes including its epitopes which may call for vaccine modifications.

## Introduction

The current COVID-19 pandemic is fueled by a more infectious emergent Omicron variant (B.1.1.529), which was first reported in South Africa and quickly spread worldwide^1^. A multitude of mutations (more than 30) in the spike gene of Omicron variant were detected, which when compared to the Alpha and Delta variants (typically less than 15)^2^, raised concerns of enhanced infectivity and immune escape potential^3,4^. Omicron variants is divided into three lineages (BA.1, BA.2, and BA.3) and was classified as the fifth variant of concern (VOC) by the World Health Organization on November 26, 2021. It has been circulating in more than 170 countries/territories.

Mutations in the SARS-CoV-2 spike gene have altered protein binding efficiency and immunogenicity, and resulted in more invasive and adaptive variants^4–9^. Previous research on Alpha (B.1.1.7) and Delta (B.1.617.2 and AY.x) variants with spike gene mutations confirmed these effects on enhancing virus transmission^4–8^. Meanwhile, as a critical antigenic recognition site, the spike protein is also the principal vaccine design target, and these observed mutations have focused attention on this modified antigen and its putative immune escape potential and antibody resistance^3,10–12^.

Ongoing concerns of emergent variants includes possible recombinants resulting from different variants replicating simultaneously in a host. Such variants, e.g., “Demicron” or “Deltacron” are controversial that if they are real recombinants or a possible sequencing error^13^.

Genome recombination is an important evolutionary mechanism for the emergence and re-emergence of human pathogens and a major source of viral evolution, for example, the well-studied “model organism” adenovirus ^14–20^, and also in coronaviruses^21–23^. Recombination accelerates virus evolution through gene(s) and “function” transference and accumulation of selective and advantageous mutations, resulting in phenotype changes that include changes in pathogenicity profiles, host species virulence, zoonotic and anthroponotic transmission, and host adaptation ^14–21,24,25^.

Although recombination events among SARS-CoV-1 and MERS-CoV were well-documented^21–23^, it has been difficult to detect the recombination signatures in SARS-CoV-2 variants due to the high degree of sequence similarity amongst SARS-CoV-2 isolates and the incomplete coverage of coronaviruses from other hosts, including pangolin^26,27^.

Previous research distinguished active recombination events among the SARS-CoV-2 nucleoprotein and ORF1ab genes by using a phylogenetic network strategy based on single nucleotide substitution or SARS-CoV-2 lineage designation^27,28^. More than thirty amino acid mutations have been identified within Omicron spike protein, some of which are shared with other variants^1^. In this study, we demonstrate that the emerging and circulating Omicron subvariants originate in part through recombination with other variants. We first investigated the spike diversity of the Omicron variants along with the shared spike mutations between Omicron and other variants of concern (VOCs) and variants of interest (VOIs). The Omicron spike amino acid sequences archived during the early transmission phase, and released in the GISAID database (submitted before January 15^th^, 2022) were accessed, include 52,563 high quality Omicron spike sequences (representing 49,609 BA.1 and 2,954 BA.2 sequences). In this study, these were analyzed with Pymol 2.0, TBTools, BioEdit, BioAider, and jvenn^29–35^. The whole genome phylogenetic trees were constructed and annotated using NextClade^36^.

### Tracking the common mutations among Omicron (BA.1 and BA.2) and variants of concern (VOCs)

Circulating Omicron variant consists of two main subvariants, BA.1 and BA.2. BA.1 subvariant was more frequently detected than BA.2 during the early transmission phase. However, BA.2 is replacing BA.1 as the dominant epidemic subvariant in more and more countries over time^37^.

Through scanning 52,563 high-quality completed Omicron spike gene sequences, most Omicron spike mutations appear stable (frequency >99%). Eighteen core mutations (frequency >99%) of BA.1 subvariant exist in NTD (A67V, del69-70, T95I, G142D, del143-145), SD (underpinning subdomain) near the S1/S2 cleavage site (T547K, D614G, H655Y, N679K, P681H), and S2 (D796Y, N856K, Q954H, N969K, L981F)(Table1).

**Table 1.**
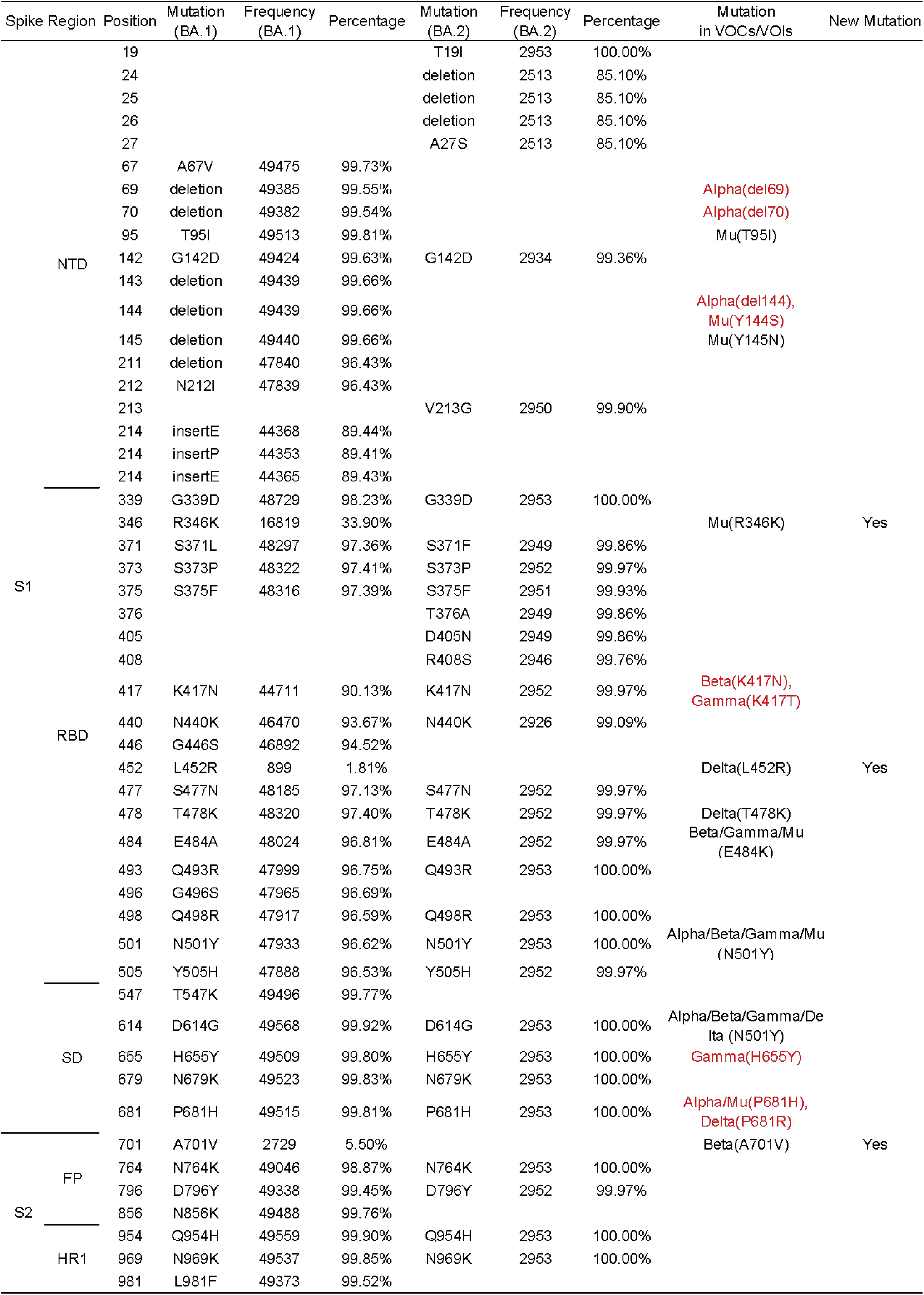
Comparison of Spike protein amino acid mutations between the Omicron subvariants and other VOCs and VOIs. 52,563 high quality Omicron spike gene sequences (49,609 BA.1 sequences, and 2,954 BA.2 sequences) released before January 15, 2022 were analyzed. The mutations that have appeared in more than 800 sequences were used in this analysis. VOCs are variants of concern; VOIs are variants of interest.

BA.1 subvariant shares nine common amino acid mutations (del69-70, delY144, K417N, T478K, N501Y, D614G, H655Y, and P681H) in the spike protein with most VOCs, suggesting a possible origin of Omicron from these VOCs. Among these shared mutations, six common ones were found in Alpha variant (del69-70, delY144, N501Y, D614G, and P681H), to which the mutations of del69-70, delY144 and P681H are exclusive; three mutations were found in Beta variant (K417N, N501Y, and D614G), to which the mutation K417N is exclusive; three mutations found in Gamma (N501Y, D614G, and H655Y), to which the mutation H655Y is exclusive; two mutations found in Delta (T478K and D614G), to which the mutation T478K is exclusive (Fig.1A and Table 1). The seven Omicron mutations exclusive to other four VOCs suggested a possible recombination origin of Omicron.

**Figure 1.**
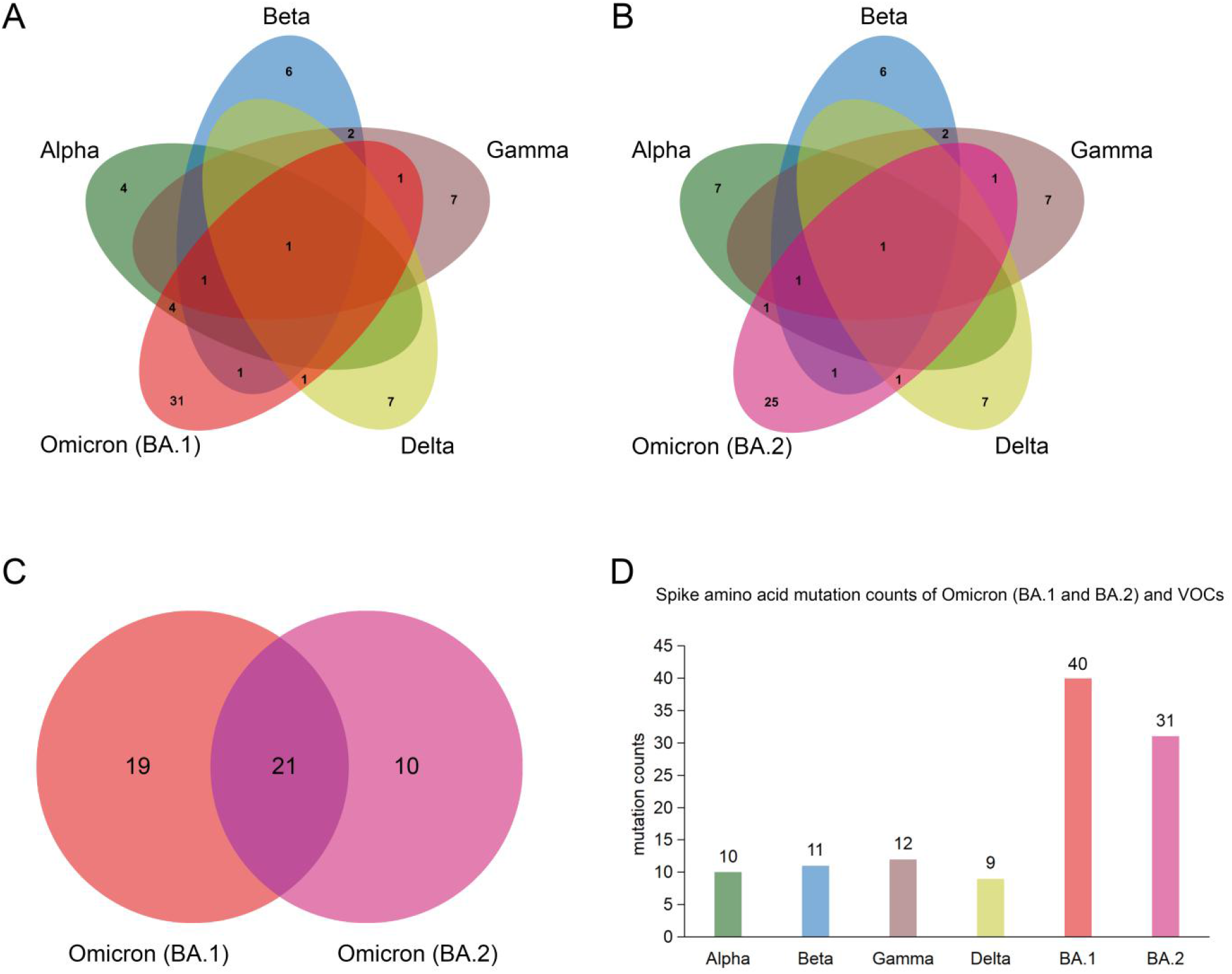
Spike protein amino acid mutations of the Omicron subvariants (BA.1 and BA.2) compared with mutations from the other four variants of concern (VOCs). **(A)** Venn diagram noting mutations of Omicron (BA.1) and those of VOCs. **(B)** Venn diagram of Omicron (BA.2) mutations compared to ones of VOCs. **(C)** Venn diagram of mutations between Omicron (BA.1) and Omicron (BA.2). **(D)** Spike protein amino acid mutation counts of Omicron (BA.1 and BA.2) subvariants compared with mutations of VOCs.

Compared to BA.1 subvariant, BA.2 shares only six amino acid mutations (K417N, T478K, N501Y, D614G, H655Y, P681H) in the spike protein with most VOCs. Among these shared mutations, three mutations were found in Alpha variants (N501Y, D614G, P681H); there were no del69-70 and delY144 mutations. The other three mutations in Beta, three mutations in Gamma, and two mutations in Delta were identical in the BA.2 and BA.1 genomes (Fig.1 B and Table 1).

BA.1 and BA.2 subvariants share twenty-one spike amino acid mutations: One in the N-terminal domain (NTD) (G142D), twelve in the receptor binding domain (RBD) (G339D, S373P, S375F, K417N, N440K, S477N, T478K, E484A, Q493R, Q498R, N501Y, Y505H), four in SD (D614G, H655Y, N679K, P681H), and four in S2 (N764K, D796Y, Q954H, N969K) (Fig.1 C and Table 1). In contrast to BA.2 subvariant, BA.1 share three additional amino acid deletions (del69-70, delY144) with the Alpha variants, suggesting a closer relationship between the BA.1 and Alpha variants (Fig.1A and 1B, and Table 1). As a whole, Omicron subvariants have a high number of amino acid mutations in the spike gene (40 in BA.1, and 31 in BA.2), of which some were found in other VOCs: Alpha (10x), Beta (11x), Gamma (12x), and Delta (9x). These mutations mainly occur in NTD and RBD (Fig.1D and Table 1).

### Tracking novel mutations and mutations with decreased frequency in the spike gene of Omicron BA.1 and BA.2

We investigated additional mutations among recently emerged BA.1 isolates and identified eight novel mutations in Omicron variant which were also found in other VOCs and VOIs. For example, mutations R346K (33.90% of 49,609 BA.1 sequences) was found in Mu variants; A701V (5.50%) was found in Beta variants; L5F (0.37%) was found in Iota variants; and T76I (0.10%) was found in Lambda variants. Most notably, multiple representative amino acid mutations in the Delta spike protein were also identified in the recently emerged Omicron subvariants (del156-167, R158G, L452R, and P681R, at percentages of 0.14%, 0.14%, 1.81%, and 0.12%, respectively. This implied possible recombination events between the Omicron and Delta strains during the pandemic. The other newly noted mutations (L141F, F643L, I1081V, S1147L, and P1162S) may have originated independently (Table 2).

**Table 2.**
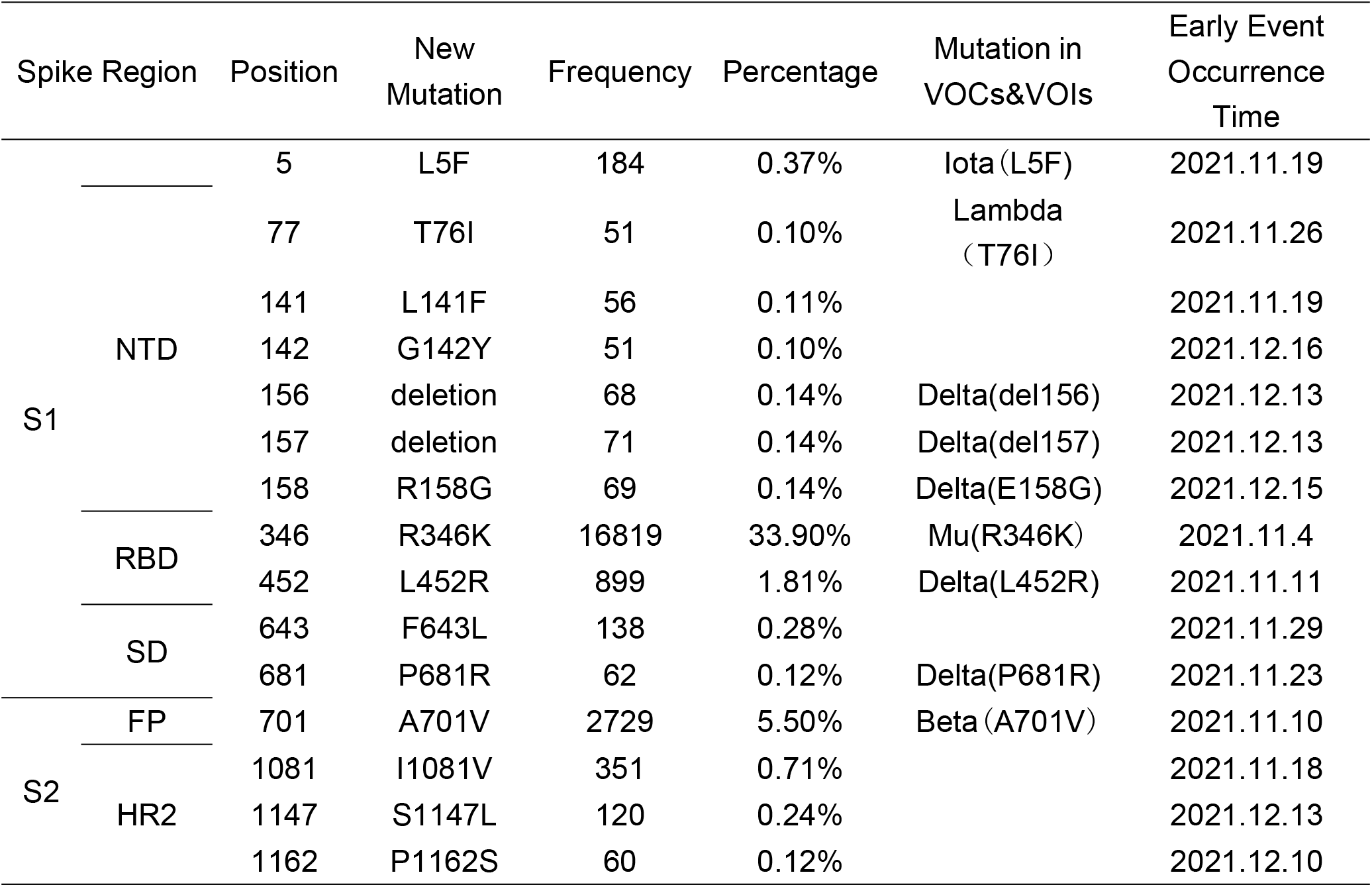
Novel mutations identified in the spike protein of the recently emerged Omicron subvariants (Released before January 15, 2022; frequency >50 sequences).

Several novel mutations were reported to be related to spike protein function, resulting in an enhancement of virus infectivity or in viral immune escape. Mutations that occurred in RBD, e.g., R346K, could result in a relatively weakened neutralizing antibody effect^3^. A L452R mutation may provide evasion from cellular immunity and increased infectivity^5,6^. The P681R as well as F643L and A701V mutations, near the S1/S2 cleavage site, may be associated with enhanced fusogenicity and pathogenicity of SARS-CoV-2 Delta variants^8^. Additionally, mutations T76I, L141F, G142Y, 156-167deletion, and R158G, located in the NTD region, were noted to affect antibody binding efficiencies and contribute to immune escape^38^. These mutations sites are mapped and shown in Fig. 2.

**Figure 2.**
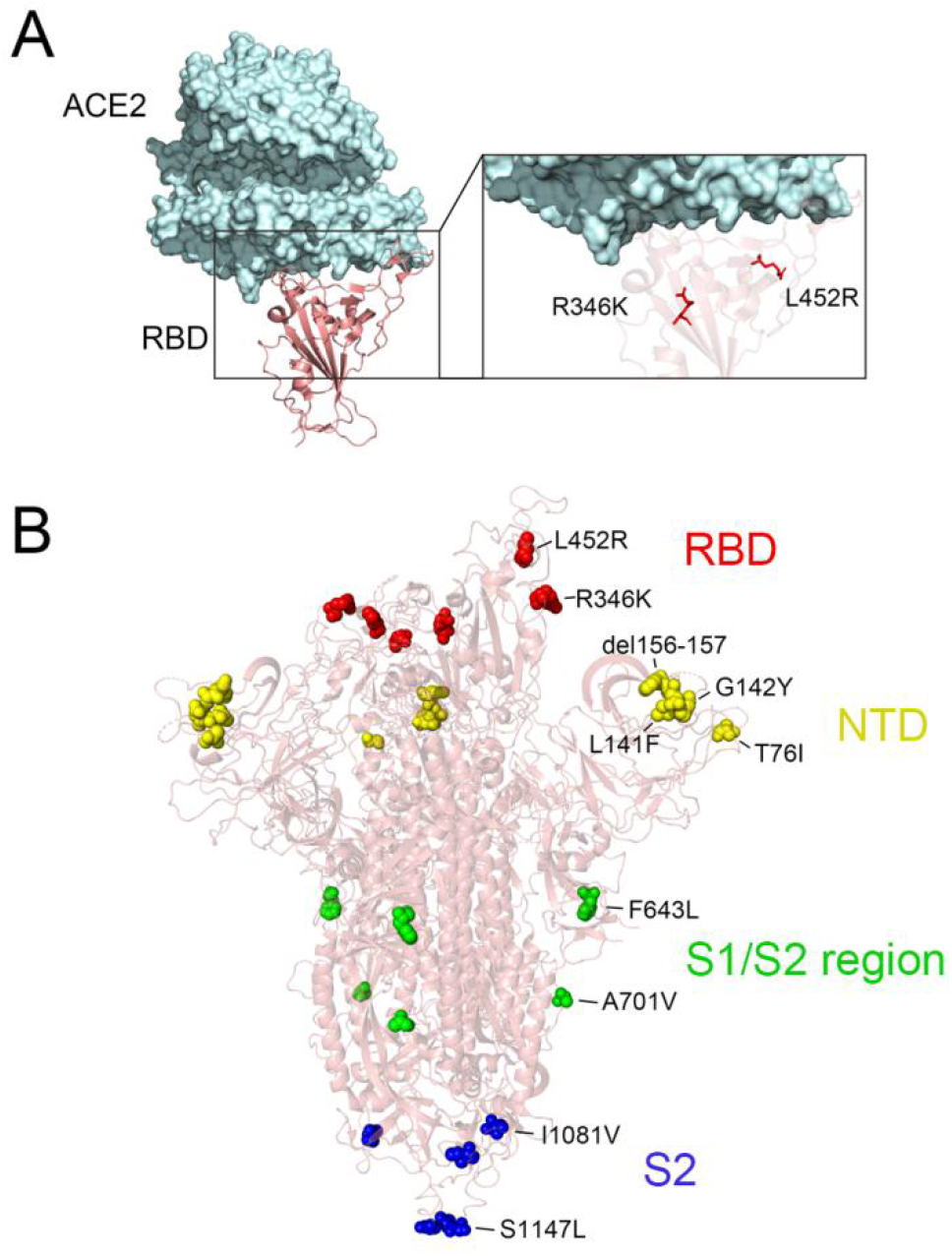
Structure of the Spike protein with amino acid mutations detected in Omicron BA.1 subvariant. **(A)** Structure of human ACE2 receptor complexed with SARS-CoV-2 Omicron RBD, mapped with the recent mutations. (**B)** Structure of SARS-CoV-2 Omicron spike protein mapped with the novel mutations. Mutated residues in each domain of the spike protein are annotated in color (red: RBD; yellow: NTD; green: S1/S2; blue: S2) using with Pymol 2.0 software through SARS-CoV-2 Omicron model PDB:7WBL and 7QO7 (Han, P. et al. Receptor binding and complex structures of human ACE2 to spike RBD from omicron and delta SARS-CoV-2. Cell, 2022, doi:10.1016/j.cell.2022.01.001).

Apparent revertant mutations are found in some dominant mutations (frequency >95%) in the BA.1 subvariant during the pandemic. Examples are the mutations in NTD (del211 and N212I) and RBD (G339D, S371L, S373P, S375F, K417N, 440K, G446S, S477N, T478K, E484A, Q493R, G496S, Q498R, N501Y, and Y505H). The frequency of insertions of the amino acids EPE at site 214 in BA.1 decreased during the pandemic from more than 95% on December 1^st^, 2021 to 89% on January 15^th^ 2022. However, BA.2 spike protein remained constant (frequency >99%), with the exception of the three amino acid deletion (LPP) found at amino acids 24-26, which decreased from more than 95% frequency on December 1^st^, 2021 to 85% on January 15^th^ 2022 (Table 1). This may possibly be due to selection pressure on the circulating Omicron strains.

### The phylogenetic network of Omicron spike genes shows novel recombination events during the pandemic

Further investigation of the Omicron subvariants examined spike protein haplotypes. These were identified, screened, and calculated with the R package tidyfst, aligned with MAFFT^39,40^. These haplotypes were analyzed with DnaSP 6.0^41^, and a subsequent phylogenetic network was constructed using PopART (http://popart.otago.ac.nz/index.shtml), with mutations annotated with Nextclade (https://clades.nextstrain.org).

The spike gene of Omicron subvariants consists of 49 representative haplotypes (each occurring in more than 50 sequences). BA.1, BA.2, R346K, L452R, A701V, and a revertant type were identified in the phylogenetic network analysis (Fig. 3A). A large number of BA.1 spike mutations delineated haplotype 2, R346K, L452R, and A701V clusters and formed distinct subgroups (detailed mutations defining each haplotype are listed in Supplementary Table 1).

**Figure 3.**
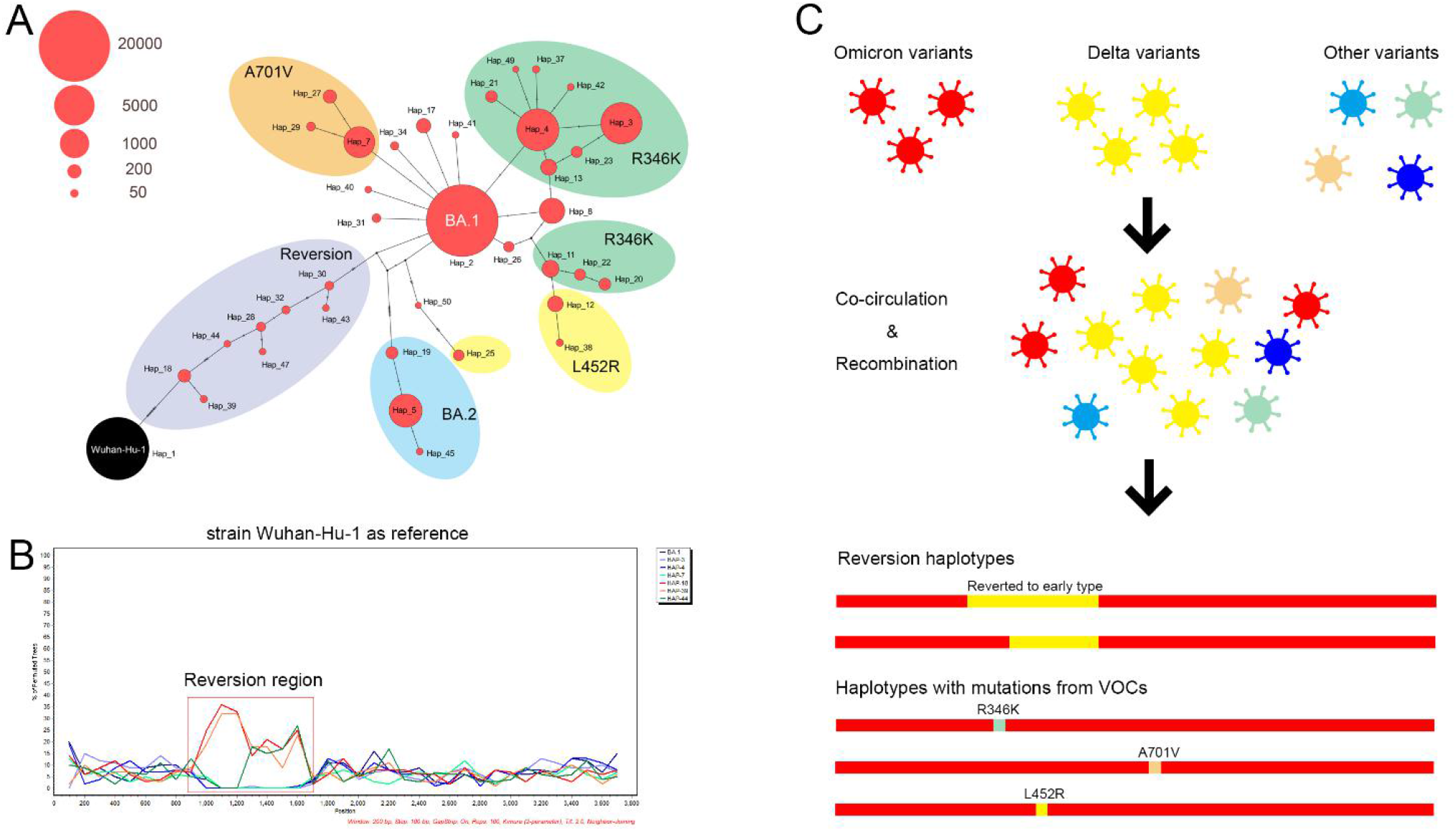
Phylogenetic network and scanning of the spike gene from representative Omicron subvariant sequences. **(A)** Representative Omicron spike protein haplotypes (each consisted of at least 50 sequences) were constructed with PopART using the median-joining method^42^. Nucleotide changes were notated with lines. The spike gene from Wuhan-Hu-1 strain was set as the root. The number of sequences in each haplotype were modified into different orders of magnitude, and subgroups based on the mutation types were delineated by color. **(B)** BootScan analysis of revertant and representative haplotypes of Omicron spike gene. Representative spike Omicron haplotypes (Hap_3, Hap_4, Hap_7) sequences and selected reversion haplotypes (Hap_18, Hap_39, Hap_44) sequences are included. Bootscan map was constructed by Simplot 3.5.1 (http://www.welch.jhu.edu/~sray/download) using neighboring-joining method with 100 bootstrap replicates.Wuhan-Hu-1 spike sequences was set as reference, reversion region was annotated. **(C)** Overview of possible evolution mechanism of reversion haplotypes and haplotypes with mutations from Delta and other variants.

Multiple nucleotide mutations were detected in the haplotypes compared with BA.1, e.g., haplotype 19 and the revertant subgroup (Hap 30, 32, 43, *etc*.) and BA.2. The L452R subgroup consists of different haplotypes with multiple nucleotide substitutions, indicating a possibly separate origin of L452R haplotypes or prior recombination events. The revertant subgroup consisted of Omicron haplotypes in which several BA.1 representative mutations were lost and appeared to have reverted to the bases of the Wu-hu-1 strain. Multiple nucleotide differences in other haplotypes occurred, likely as multiple independent mutation events, or perhaps as recombination events among highly similar sequences. Haplotype 25 in L452R subgroup, with multiple nucleotide differences compared with BA.1, could have resulted through recombination between Omicron and Delta variants, gaining the mutation L452R from Delta and losing multiple mutations from Omicron (Fig. 3A). Some of these “Demicron” or “Deltacron” haplotypes are being tracked by the UK Health Security Agency (https://www.gov.uk/government/publications/sars-cov-2-variants-of-public-health-interest/sars-cov-2-variants-of-public-health-interest-25-february-2022) and underway to confirm by Santé publique France (https://t.co/tVAKmHRYSy). Bootscan analysis of Omicron spike sequences also indicated that the reversion haplotypes (Hap_18, Hap_39, Hap_44) were more similar to Delta variants when compared to typical Omicron haplotypes (Fig. 3A and 3B).

Furthermore, single nucleotide differences could also originate from recombination events among highly similar strains. Loops detected in phylogenetic networks also indicate possible recombination events among highly similar Omicron variants or subvariants (Fig. 3A and 3C). Multiple newly detected or recent mutations in the Omicron spike gene make it possible to trace a putative mutation origin from representative mutations in VOIs or VOCs, especially the Delta variant, which suggests possible recombination events between Omicron and Delta variants (Table 2).

### Co-infections of different SARS-CoV-2 variants in the population accelerates their evolution through recombination

Virus co-infection and recombination can amplify pathogenicity, for example, the well-studied “model organism” adenovirus ^14–20^, and also in coronaviruses^21–23^. SARS-CoV-2 has been shown to co-infect and recombine ^26,43^. In host populations with disproportionate immunocompromised conditions, such as Africa^44^, the possibility of long-term infections of SARS-CoV-2 variants may be higher than in populations otherwise healthy and/or vaccinated. A case report described prolonged infectious SARS-CoV-2 shedding up to 70 days from an asymptomatic immunocompromised individual with cancer^43^. A SARS-CoV-2 isolated from her presented with four new mutations within the spike protein and also eight in structural proteins and polymerase region. The marked within-host genomic evolution of SARS-CoV-2 with continuous turnover of dominant viral variants was observed^43^. Under reduced immune pressure or immune-suppression, long-term infections create conditions and increase the likelihood of simultaneous co-infections with multiple SARS-CoV-2 variants, and optimizing conditions for genome recombination. For example, on June 10, 2021, a passenger on a flight from Johannesburg, South Africa to Shenzhen, China tested positive for SARS-CoV-2^26^. The patient was found to be coinfected with two SARS-CoV-2 variants: Beta and Delta, with the ratio of the relative abundance between the two variants maintained at 1:9 (Beta: Delta) in a 14-day period. Furthermore, putative evidence of recombination in the Orf1ab and spike genes was shown^26^. Such recombination events may not be rare, especially considering that there are hundreds of variants circulating in the general population.

Among the Omicron subvariants and VOCs, many shared mutations were identified in this study. We speculate that some of the Omicron spike protein mutations resulted from co-infections of variants. Recombination among diverse variants may have contributed to the shared presence of different mutations between the VOCs. For example, the BA.1 subvariant has three more Alpha-related mutations (del69-70, delY144) than BA.2, and therefore may be phylogenetically closer to the Alpha variant, suggesting that Alpha or other unknown variants that carry these mutations may have contributed to the emergence of the BA.1 subvariant (Table 1). Multiple mutation differences causing reversion haplotypes may have originated from the recombination between the Omicron and other variants (Fig. 3 and Supplementary Table 1).

Except for shared mutations, many other mutations (30 in BA.1 and 25 in BA.2) could not be accounted for among previous dominant variants (Fig. 1). Because the Omicron variants are believed to have emerged in South Africa^1^, we speculate that some of these spike mutations may have been produced by long-term virus infections in immunocompromised patients. It was previously reported that evolution of SARS-CoV-2 in an immunosuppressed COVID-19 patient led to immune escape variants^45,46^. Deletions in NTD, for example, delY144, were detected in multiple immunosuppressed COVID-19 patients, which resulted in immune escape^45–47^.

### This extended pandemic is likely yielding novel recombinant SARS-CoV-2 variants

A recently reported recombinant SARS-CoV-2, “Deltacron” or “Demicron”, and its genome sequences, elicited controversy and concerns of sequencing errors and sample contamination^13^. Nevertheless, it was confirmed that co-infections by Omicron and Delta variants have already occurred in specific populations (https://www.gov.uk/government/publications/sars-cov-2-variants-of-public-health-interest/sars-cov-2-variants-of-public-health-interest-11-february-2022). Recombination among the extant variants may lead to the emergence of new variants. A total of 10 cases of “Deltacron” are underway to confirm by Santé publique France (SPF) (https://t.co/tVAKmHRYSy). In our study, multiple VOC and VOI mutations were detected in Omicron variants circulating before January 15, 2022 (Fig. 1 and Table 1). The integration of these mutations may lead to changes in phenotype. Five additional typical amino acid mutations in Delta variants were also identified in recently emergent Omicron isolates (before January 15, 2022) (Table 2). For example, 899 Omicron sequences of high quality contained L452R mutation reported for the Delta variant (Fig. 4A). Whole genome analysis also corroborated the diversity among these L452R containing Omicron genomes. The mutation profiles among whole genomes of BA.1 are diverse, and the sequences branched to diverse clades by phylogenetic analyses (Fig. 4B).

**Figure 4.**
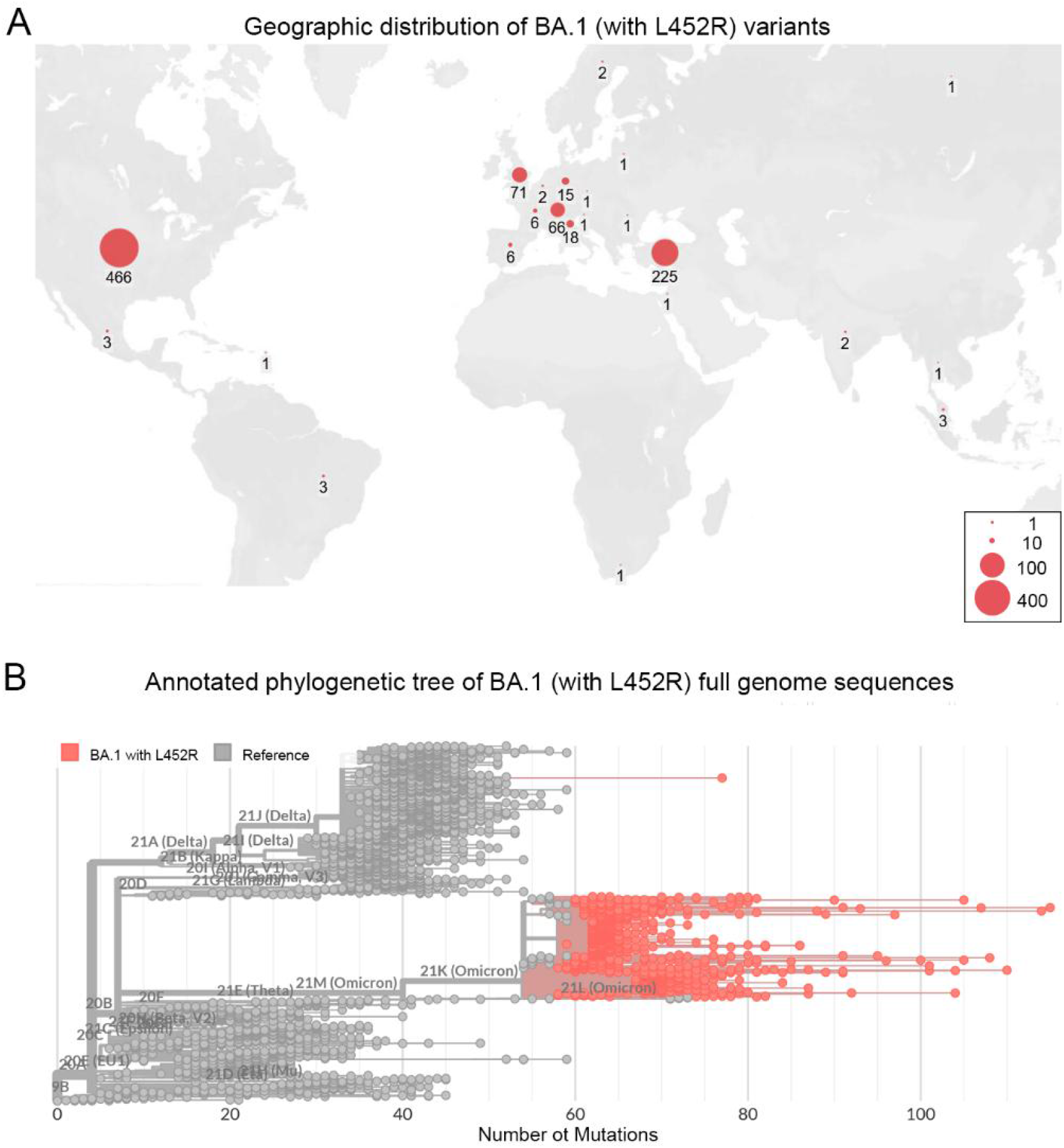
Geographic distribution and whole genome analyses of BA.1 (with L452R) variants. **(A)** Geographic distribution of BA.1 (with L452R) subvariants, with the number of genome sequences noted. (**B)** Whole genome phylogenetic tree highlighting the BA.1 (with L452R) subvariant, clades were annotated using NextClade (Hadfield et al., 2018). Low quality sequences were excluded. 891 SARS-CoV-2 BA.1 with L452R spike amino mutation full genome sequences submitted to the GISAID database before January 15^th^, 2022 and reference sequences from SARS-CoV-2 each clade were included. The red circles are BA.1 variants.

## Conclusion

By analyzing sequences from a large number of Omicron subvariants, we identified diverse recombination events between two Omicron subvariants and several SARS-CoV-2 variants, suggesting that co-infection and subsequent genome recombination play important roles in the on-going evolution of SARS-CoV-2. Some of the recombination events may have led to modifications in protein function and viral fitness. Continued monitoring of SARS-CoV-2 genomes for mutations is critically important to our understanding of its evolution and impact on human health, and is also essential for the recognition of changes to viral epitopes that would require vaccine modifications.

## Author contribution statement

JO and QZ contribute to study design and manuscript writing. JO, WL, XW, TZ, BD, PY, YR, LQ, and QZ contribute to data analysis and data visualization. WZ, DS, JC, JW and QZ contribute to manuscript revision.

## Acknowledgments

We gratefully acknowledge the authors, originating and submitting laboratories of the sequences from GISAID’s EpiFlu™ Database on which this research is based. All submitters of data may be contacted directly via www.gisaid.org.

## Funding statement

This work was supported by grants from the National Key Research and Development Program of China (2018YFE0204503), National Natural Science Foundation of China (32170139 and 81730061), and Natural Science Foundation of Guangdong Province (2018B030312010 and 2021A1515010788).

## Competing interests

The authors declare no competing interests.

## Supplementary information

**Supplementary Table 1**. The annotation of haplotypes of Omicron spike protein.

**This version was submitted for peer review on March 3, 2022**.

## Notes

### Competing Interest Statement

The authors have declared no competing interest.

